# A novel strategy for single-cell metabolic analysis highlights dynamic changes in immune subpopulations

**DOI:** 10.1101/2020.01.21.914663

**Authors:** Patricia J. Ahl, Richard A. Hopkins, Wen Wei Xiang, Bijin Au, Nivashini Kaliaperumal, Anna-Marie Fairhurst, John E. Connolly

## Abstract

A complex interaction of anabolic and catabolic metabolism underpins the ability of leukocytes to mount an immune response. Their capacity to respond and adapt to changing environments by metabolic reprogramming is crucial to their effector function. However, current methods lack the ability to interrogate this network of metabolic pathways at the single cell level within a heterogeneous population. Here we present Met-Flow, a novel flow cytometry-based method that captures the metabolic state of immune cells by targeting key proteins and rate-limiting enzymes across multiple pathways. We demonstrate the ability to simultaneously measure divergent metabolic profiles and dynamic remodeling in human peripheral blood mononuclear cells. Using Met-Flow, we discovered that glucose restriction and metabolic remodeling drive the expansion of an inflammatory central memory T cell subset. This method captures the complex metabolic state of any cell as it relates to its phenotype and function, leading to a greater understanding of the role of metabolic heterogeneity in immune responses.

## Introduction

The immune status of a given cell type is defined by its underlying metabolic state. Leukocytes utilize metabolic pathways to coordinate immune specific gene expression at the epigenetic, transcriptional, post-transcriptional and post-translational levels. In T cells, glycolysis plays an important role in effector function and cytokine production^1^, and high activity through the AKT signaling pathway during activation supports both increased glycolysis and oxidative phosphorylation (OXPHOS) of naïve T cells^2^. In the context of activation in antigen presenting cells (APC), glycolysis, glycogen metabolism and fatty acid synthesis are required for immuno-stimulatory function ^3–6^. Conversely, the formation of regulatory T cell subsets (Treg) requires fatty acid synthesis ^7^, whereas tolerogenic dendritic cells require fatty acid oxidation for active suppression ^8,9^. This metabolic switch to lipid metabolism is driven by increased signaling of the mechanistic target of rapamycin (mTOR) pathway, measured by flow cytometry of phosphorylated proteins (Phos-Flow)^10,11^. These findings illustrate the critical role of multiple metabolic pathways in shaping cellular phenotype and function.

Multiplexing the metabolic state of cells and immune function is limited by available technologies. The field of immunology is dominated by high dimensional single cell analysis using flow cytometry, mass cytometry, and single cell RNA sequencing (scRNAseq), whereas bulk cellular analysis technology is often used to capture metabolic respiration. However, these technologies are largely incompatible with analysis of heterogeneous cellular populations at a protein level.

Here we present Met-Flow, a high-parameter flow cytometry method utilizing antibodies against metabolic proteins that are critical and rate-limiting in their representative pathways. The cell’s potential to flux through anabolic pathways was examined by the measurement of fatty acid synthesis and an arginine metabolism associated protein. The analysis of catabolic pathways encompassed the quantification of proteins involved in glycolysis, the pentose phosphate pathway (PPP), the tricarboxylic acid (TCA) cycle, oxidative phosphorylation (OXPHOS) and fatty acid oxidation. The capacity for phosphate and glucose uptake was measured by the expression level of metabolic transporters, as well as an antioxidant enzyme that affects oxidative stress (Supplementary Table 1).

The protein composition of these rate-limiting enzymes defines the cellular capacity of a metabolic pathway. Furthermore, dynamic cellular differentiation engages rapid post-transcriptional and post-translational mechanisms, thus affecting the concentrations of metabolic pathway-associated proteins. Met-Flow allows us to simultaneously capture the state of key metabolic pathways on a single-cell, protein level, thus overcoming the inherent drawbacks of metabolic mRNA analysis, including: the temporal discord between mRNA abundance with protein concentration^12^. Moreover, dynamic cellular differentiation engages rapid post-transcriptional and post-translational mechanisms, that are not regulated by gene expression^13^. Combined, these limitations highlight the importance of protein-level analysis.

Here, we demonstrate the ability of Met-Flow to measure divergent metabolic states across healthy human peripheral blood mononuclear cells (PBMCs) and draw novel associations between the metabolic profile of a cell with its subset phenotype, activation status, and immunological function. With the ability to capture metabolic heterogeneity on a single cell level, Met-Flow provides important insights into the understanding of the metabolic state across any cell type.

## Methods

### Peripheral Blood Mononuclear Cell (PBMC) isolation

PBMC were isolated from cone blood of healthy donors (IRB NHG DSRB 2000/00828) using ficoll density gradient centrifugation (Ficoll-Paque, GE Healthcare). Whole blood was diluted in a 1:1 ratio with PBS (Gibco^TM^, ThermoFisher, 10010023) supplemented with 2 mM EDTA (PBS-EDTA) (Invitrogen^TM^, ThermoFisher, 16676038). The diluted blood was layered on top of the ficoll in a 2:1 diluted blood to ficoll ratio. The sample was spun at 400 g for 30 min without brake 21 °C. After centrifugation, the PBMC layer was carefully removed and washed twice in PBS-EDTA. Cells were frozen down in freezing medium containing FBS (Hyclone, GE Healthcare, SH30071.03) and 10 % dimethyl sulfoxide (DMSO) at 50×10^6^ PBMC/ml overnight at −80 °C and subsequently stored in liquid nitrogen.

### PBMC and T cell culture

PBMCs were thawed in a 37 °C water bath and washed with 10 ml of complete (c)RPMI containing RPMI 1640 (Gibco^TM^, ThermoFisher, 11875093), 10 % FBS, 100 U/ml penicillin and 100 μg/ml streptomycin (Gibco^TM^, ThermoFisher, 15140122), 1 mM sodium pyruvate (Gibco^TM^, ThermoFisher, 11360070), 2 mM L-glutamine (Gibco^TM^, ThermoFisher, 35050061), 1X nonessential amino acids (Gibco^TM^, ThermoFisher, 11140050), 15 mM HEPES (Gibco^TM^, ThermoFisher, 15630080). T cells were isolated from PBMCs by three sequential rounds of magnetic separation using CD3 Microbeads (Miltenyi Biotec, 130-050-101), according to the manufacturer’s instructions. PBMC and T cells were seeded at 1×10^6^ PBMC in a 96-well flat-bottom plate and rested in cRPMI for 1 hour. After resting, cells were stimulated for 24 h with Gibco™ Dynabeads™ Human T-Activator CD3/CD28 (ThermoFisher, 11131D) in a bead-to-cell ratio of 0.5:1 in simultaneous presence or absence of 2 mM 2-Fluoro-2-deoxy-D-glucose (2-FDG) (Sigma, F5006) at 37°C, 5% CO_2_.

### Flow cytometry staining

Ten metabolic proteins were chosen and optimized based on their critical role in specific metabolic pathways (Table 1). The purified metabolic antibodies were purchased from Abcam and custom conjugated by Becton Dickinson (BD) using their fluorochromes, unless otherwise indicated; SLC20A1 (clone EPR11427(2), BD AF647, Abcam ab231703), ACAC (clone EPR4971, BD BUV496, Abcam ab231686), HK1 (clone EPR10134(B), BD BUV661, Abcam ab234112), CPT1A (clone 8F6AE9, BD V450, Abcam ab231704), IDH2 (clone EPR7577, BD BB790, Abcam ab231695), G6PD (clone EPR6292, BD BUV395, Abcam ab231690), GLUT1 (clone EPR3915, Abcam AF488, ab195359), ASS1 (clone EPR12398, BD AF700, Abcam ab231684), PRDX2 (clone EPR5154, BD BUV615, Abcam ab231702), ATP5A (clone EPR13030(B), Abcam AF594, ab216385). These metabolic proteins are differentially localized to the mitochondria, the cell surface or the cytosol (Table 1). In addition, antibodies to surface and intracellular markers were used to phenotype 11 leukocyte subsets in PBMCs to generate a 27 color flow cytometry panel; CD4 (clone SK3, BD, BV480, 566104), CD8 (clone SK1, BD, BUV805, 564912), and CD3 (clone UCTH1, BD, BB630, 624294) for T cells; HLA-DR (clone G46-6, BD, BV786, 564041), CD11c (clone B-ly6, BD, BB700, 624381), for myeloid and CD123 (clone 9F5, BD, BV650, 740588) for plasmacytoid dendritic cells, IgM (clone G20-127, BD, BUV805, 624287), IgD (clone IA6-2, BD, BV480, 566138) and CD19 (clone HIB19, BD, BB660, 624295) for B cells; CD16 (clone 3G8, BD, BV750, 624380) and CD14 (clone M5E2, BD, PE-Cy7, 557742) for Monocytes; and NK subsets using CD56 (clone NCAM16.2, BD, PE-Cy5, 624350), as well as CD45 (clone 2D1, BD, BUV563, 624284), PD-1 (clone MIH4, BD, PE, 557946), ILT3 (clone ZM3.8, BD, BV605, 742807), CD69 (clone FN50, BD, APC-H7, 560737), CD86 (clone 2331/FUN-1, BD, BUV737, 564428) and live-dead dye FVS575V (BD, BV570, 565694). The modified T cell panel included CCR7 (clone G043H7, Biolegend, BV650, 353134), CD45RA (cloneHI100, BD, PE, 561883), CD25 (clone 2A3, BD, PE-Cy7, 335789), FOXP3 (clone PCH101, eBioscienceTM, Thermo Fisher, PE-Cyanine5.5, 35-4776-42) and CD14 (M5E2, BD, BV570, 624298) for Monocytes.

PBMCs or purified T cells were stained for 30 minutes on ice with the antibodies specific for extracellular proteins in Brilliant Stain Buffer (BD, 563794). Following incubation, cells were washed with cold PBS and centrifuged at 300 g, 5 minutes, 3 times. Cells were fixed and permeabilized using eBioscience Foxp3/Transcription Factor Staining Buffer Set (Invitrogen, Catalog Number 00-5523-00) according to manufacturer’s instructions. We then washed the cells in PBS as previously described and stained with intracellular antibodies in permeabilization buffer for 1 hour at room temperature. Subsequently, cells were washed once in permeabilization buffer followed by a PBS wash. Samples were acquired on a X-30 FACSymphony (BD) with FACS Diva Version (BD, Version 8.0.1) software. Analysis was completed using FlowJo (BD, version 10.5.2).

### Phos-Flow staining

Purified T cells were isolated and stimulated as described above. After incubation with different treatments, cells were added to a 96-well V-bottom plate and spun down at 1500g for 1 minute at 4°C. Cells were then stained with live-dead dye FVS575V (BD, BV570, 565694) for 5 minutes and washed by adding 150 μl PBS and spinning at 3000rpm for 1 minute at 4°C. Following this, Fix Buffer I (BD, 557870) was added at 150 μl per well and incubated for 10 minutes at 37°C. After fixation and washing as described above, 150 μl of Perm/Wash Buffer I (BD, 557885) was added and incubated for 30 minutes, in the dark, at room temperature. After permeablization, cells were stained with an antibody cocktail mix for 1 hour at room temperature in Perm/Wash Buffer I, including the antibodies CD4 (clone SK3, BD, BV480, 566104), CD8 (clone SK1, BD, BUV805, 564912), CD3 (clone UCTH1, BD, BB630, 624294), HLA-DR (clone G46-6, BD, BV786, 564041), CD16 (clone 3G8, BD, BV750, 624380), CD45 (clone 2D1, BD, BUV563, 624284), CD69 (clone FN50, BD, APC-H7, 560737), CCR7 (clone G043H7, Biolegend, BV650, 353134), CD45RA (cloneHI100, BD, PE, 561883), CD25 (clone 2A3, BD, PE-Cy7, 335789), CD14 (M5E2, BD, BV570, 624298), the phosphorylated ribosomal protein S6 (Ser240/244, clone D68F8, AF647, 5044), as well as the abovementioned 9 metabolic antibodies. Finally, cells were washed and acquired on the X-30 FACSymphony.

### GM-CSF staining

Purified T cells were stimulated as previously described and GM-CSF was measured using the GM-CSF Secretion Assay Enrichment and Detection Kit (PE, Miltenyi, 130-105-760). The manufacturer’s instructions were modified to a 96-well format with final volumes of 200 μl per well. Following the GM-CSF kit protocol, cells were additionally stained with CD4 (clone SK3, BD, BV480, 566104), CD8 (clone SK1, BD, BUV805, 564912), CD3 (clone UCTH1, BD, BB630, 624294), HLA-DR (clone G46-6, BD, BV786, 564041), CD16 (clone 3G8, BD, BV750, 624380), CD45 (clone 2D1, BD, BUV563, 624284), CD69 (clone FN50, BD, APC-H7, 560737), CCR7 (clone G043H7, Biolegend, BV650, 353134), CD45RA (cloneHI100, BD, PE, 561883), CD25 (clone 2A3, BD, PE-Cy7, 335789), CD14 (M5E2, BD, BV570, 624298), CD56 (clone NCAM16.2, BD, PE-Cy5, 624350) and live-dead dye FVS575V (BD, BV570, 565694). Subsequently, cells were fixed and permeabilized in Foxp3/Transcription Factor Staining Buffer as previously described, and stained with the abovementioned metabolic antibodies, before acquiring on the X-30 FACSymphony.

### Cytokine and chemokine analysis

Supernatants from stimulation experiments were collected and stored at −80 °C for analysis. Cytokine and chemokine profiles were analyzed using a multiplexed, bead-based kit (Milliplex 41-plex human cytokine panel 1, Millipore, MA, USA) on the FLEXMAP 3D system (Luminex Corporation, TX, USA).

### Real-time metabolic characterization using extracellular flux analysis

Glycolytic function and mitochondrial respiration were measured by extracellular acidification rate (ECAR, mpH/min) and oxygen consumption rate (OCR, pmol/min) using the XFe96 extracellular flux analyzer (Seahorse Bioscience, Massachusetts, USA). 200,000 cells per well were plated in a 96 well plate and pre-treated for 24 h in the presence or absence of CD3/28 beads and 2-FDG in cRPMI. Respiration was measured in XF Assay Modified Media with L-glutamine (2 mM), sodium pyruvate (1 mM) with or without 11 mM Glucose (Sigma-Aldrich, Merck, G8769) for OCR and ECAR measurements, respectively. To measure glycolytic parameters, the glycolytic stress test kit (Seahorse Bioscience, 103020-100) was used, containing Glucose (10 mM), oligomycin (2 μM) and 2-deoxy-glucose (50 mM). Mitochondrial respiration parameters were measured using the mitochondrial stress test kit (Seahorse Bioscience, 103015-100), by sequentially adding oligomycin (2 μM), Carbonyl cyanide-4 (trifluoromethoxy) phenylhydrazone (FCCP) (1.5 μM), Rotenone and Antimycin A (1 μM).

### High-dimensional and statistical analysis

Statistical analysis was performed using Prism (Graphpad, version 8.2.0). Data were compared using either T-tests for paired analysis or non-parametric one-way ANOVA with Dunn’s Multiple Corrections, unless otherwise stated. Data is represented as the mean ± standard deviation (SD). P values <0.05 were considered significant; where *P<0.05, **P<0.01, ***P<0.001, ****P<0.00001. High-dimensional analysis by Fast Fourier Transform-accelerated Interpolation-based t-distributed stochastic neighbor embedding (Fit-SNE) was performed using FlowJo (BD, Version 10.6.1). Heatmaps were generated using a web-enabled tool (Heatmapper ^14^). Chord plots^15^ were generated by Spearman correlation analysis of gMFI in one immune population relative to the gMFI of all other subsets. Analysis of scRNAseq data of 68k PBMCs and 5k PBMCs was done using previously published data and R studio ^16,17^.

## Results

### Immune cells exhibit divergent metabolic profiles at a protein level

Innate and adaptive immune responses are orchestrated by leukocytes, which require metabolic remodeling and mitochondrial signaling to exert their function ^18–21^. In our studies we set out to develop the capability to measure metabolic profiles across multiple immune subsets in a heterogeneous population on a single-cell, protein level.

A 27-parameter flow cytometry panel was built, which included 10 critical metabolic proteins, encompassing rate-limiting enzymes, anabolic and catabolic pathways and transporters (Supplementary Table 1, Supplementary Fig. 1a), as well as phenotypic markers to analyze 11 major leukocyte subsets. Using the FitSNE algorithm, cellular subsets from 12 donor samples were clustered into their immune phenotype with 15,000 cells per leukocyte population from each donor, based on similarities in expression profiles of individual cells ^22,23^. This methodology successfully clustered the populations based on differential expression of both the lineage and metabolic proteins (Fig. 1a, Supplementary Fig. 1b). To determine whether the immune cell subsets could be identified by their metabolic phenotype alone, the clustering analysis was performed using the expression profiles of only the 10 metabolic proteins. Using the divergent expression levels of metabolic proteins alone clustered the populations into CD3^+^ T cells, CD56^+^ NK cells, CD19^+^ B cells, HLA-DR^+^/CD11c^+^/CD14^−^ myeloid Dendritic cells (mDCs) and CD14^+^ monocytes (Fig. 1b), which were retrospectively identified by their lineage marker expression (Supplementary Fig. 1c), and confirmed by an overlay of conventionally gated immune populations (Supplementary Fig. 1d). Both monocytes and mDCs segregated into distinct, metabolically defined islands. The monocytes separated out into 2 subpopulations, mainly due to a difference in expression of the TCA cycle enzyme IDH2 (Fig. 1b). Unlike the projection of phenotypic markers that separated out the functional CD4^+^ and CD8^+^ subsets (Fig. 1a), using metabolic protein expression profiles alone showed a similar metabolic profile across all CD3^+^ T cells (Fig. 1b, Supplementary Fig. 1c,d).

**Figure 1.**
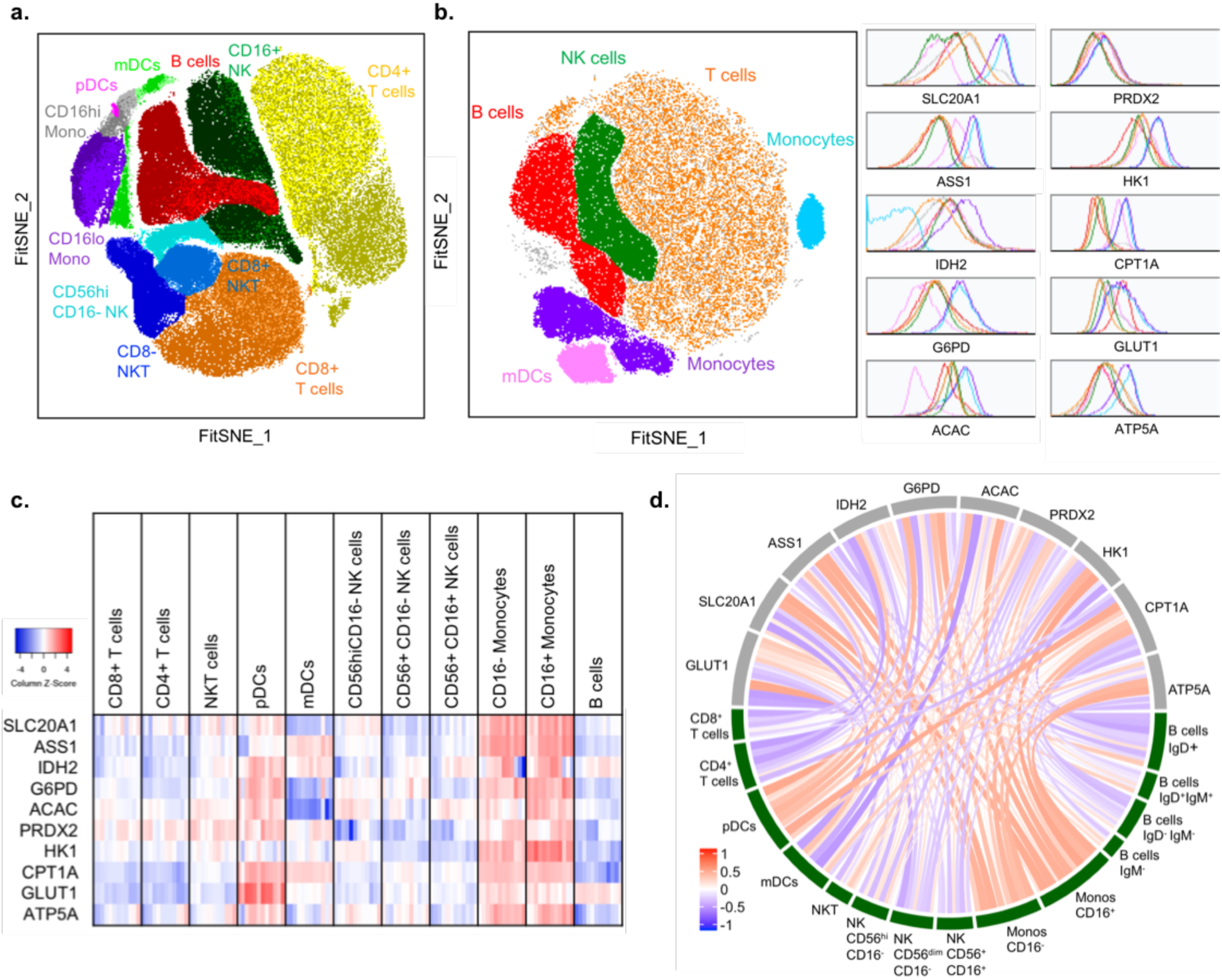
Protein level analysis shows divergent metabolic profiles in leukocytes. (a) FitSNE projection of both phenotypic and metabolic proteins, and (b) FitSNE projection of metabolic proteins only, with corresponding expression in each population, representing n=12 samples from 4 independent experiments. (c) log2 of gMFI expression of each immune cell type (n=12). (d) Chord visualization using spearman correlation between metabolic protein and immune phenotype. A positive correlation is presented in red, a negative correlation is presented in blue based on the r value (n=9).

In comparison with scRNAseq analysis of isolated PBMC populations, we showed the ability of Met-Flow to define immune cells by their metabolic state with 10 metabolic proteins, which was comparable to the resolution of >500 metabolic genes by scRNAseq^16,17^ (Supplementary Fig. 1d). Unlike the protein level analysis, the expression of the same 10 metabolic genes alone were not able to resolve immune populations at the RNA level (Supplementary Fig. 1e). There is a well-characterized contribution of both post-transcriptional and post-translational modifications that regulate metabolic genes^1,2,24^. Our data demonstrates the strong correlation of the metabolic protein profile with distinct leukocyte subsets. Furthermore, the ability to identify these subsets using transcriptome data requires a greater amount of dimensionality compared to when using the protein-based Met-Flow method, thus reducing the burden for advanced analytical techniques.

Using a comparative heatmap analysis of the geometric mean fluorescence intensity (gMFI) of each protein, we showed metabolic heterogeneity across leukocytes, where each leukocyte population was gated based on specific lineage markers (Fig. 1c, Supplementary Fig. 2a,b). In plasmacytoid DCs (pDCs), our data showed higher levels of IDH2, ATP5A, G6PD and GLUT1 reflecting heightened capacity for OXPHOS, the TCA cycle, PPP and glucose uptake compared to mDCs (Fig. 1c, Supplementary Fig. 2c). In both CD16^hi/lo^ monocyte subsets, the expression of all metabolic proteins is high in comparison to the other populations. Inflammatory CD16^+^ monocytes expressed higher G6PD, ACAC and HK1 in comparison to the CD16^−^ population (Fig. 1c, Supplementary Fig. 2d). Analysis of B cells showed significantly higher GLUT1 and IDH2 in comparison to T and NK subsets (Fig. 1c, Supplementary Fig. 2e-f). The increased GLUT1 and IDH2 indicates a high capacity for glucose uptake and OXPHOS, which has previously been shown to play a critical role for B cell activation by mTOR signaling, mitochondrial membrane potential remodeling and ROS production ^25,26^. Across CD16^−^ NK cell subsets, our data demonstrates divergent metabolic profiles of CD56 bright cells compared to the dim population (Fig. 1c, Supplementary Fig. 2g). The CD56 bright cells express higher SLC20A1, ASS1, ACAC and HK1, whereas CD56 dim cells show greater expression of CPT1A and GLUT1. Moreover, in comparison to CD4^+^ T cells, we demonstrated higher expression of IDH2, G6PD, ACAC, CPT1A, GLUT1 in NKT cells (Fig. 1c, Supplementary Fig. 2h). Lastly, GLUT1 and HK1 are expressed at similar levels between CD4^+^ and CD8^+^ T cells (Fig. 1c, Supplementary Fig. 2i), as both subsets similarly rely on glycolytic flux ^27^, however there is a significant difference in G6PD indicating a dissimilarity in capacity for flux through the PPP. Additionally, the relative correlation between immune subsets of a given phenotypic marker to each metabolic protein was measured. This data showed an increased or decreased association between specific metabolic pathways and individual leukocyte populations reflecting metabolic heterogeneity of cellular human PBMC populations (Fig. 1d).

Collectively, this data demonstrated the ability of this novel immuno-metabolic flow cytometry panel to capture differential metabolic profiles within a heterogeneous immune cell population. Met-Flow measured single cell and protein level metabolic states and provided novel correlations between immune subpopulations and specific metabolic pathways.

### Dynamic metabolic reprogramming occurs during T cell activation

With the ability to measure divergent metabolic profiles across resting immune populations, the relationship between metabolism, leukocyte activation and maturation was tested using purified T cells. To explore metabolic dynamics, beads coated with anti-CD3 and anti-CD28 (CD3/28) were added to activate T cells by TCR engagement and co-stimulatory signal ^28^. A modified flow cytometry panel was used to include T cell memory markers, with a focus on CD4^+^ T cells. Stimulation of the T-cells altered activation-dependent protein levels, with the highest fold change increase observed in CD25 expression, followed by CD69 and HLA-DR (Fig. 2a). Simultaneous measurement of metabolic protein expression showed a 3-fold induction of GLUT1, suggesting a significantly increased capacity for glucose transport in these activated cells (Fig. 2a, Supplementary Fig, 3a,b). Moreover, the analysis showed over 2-fold inductions of IDH2, ACAC, G6PD, ASS1 and PRDX2, indicating an increased capacity for flux through the TCA cycle, fatty acid synthesis, oxidative PPP, arginine synthesis and the antioxidant response pathways, respectively (Fig. 2a, Supplementary Fig. 3a,b). HK1, ATP5A and CPT1A were also significantly higher following activation, showing increased capacity for flux through glycolysis, OXPHOS and fatty acid oxidation (Fig. 2a,b). Cumulatively, the data demonstrated that differential reprogramming of multiple metabolic pathways is closely linked to T cell activation.

**Figure 2.**
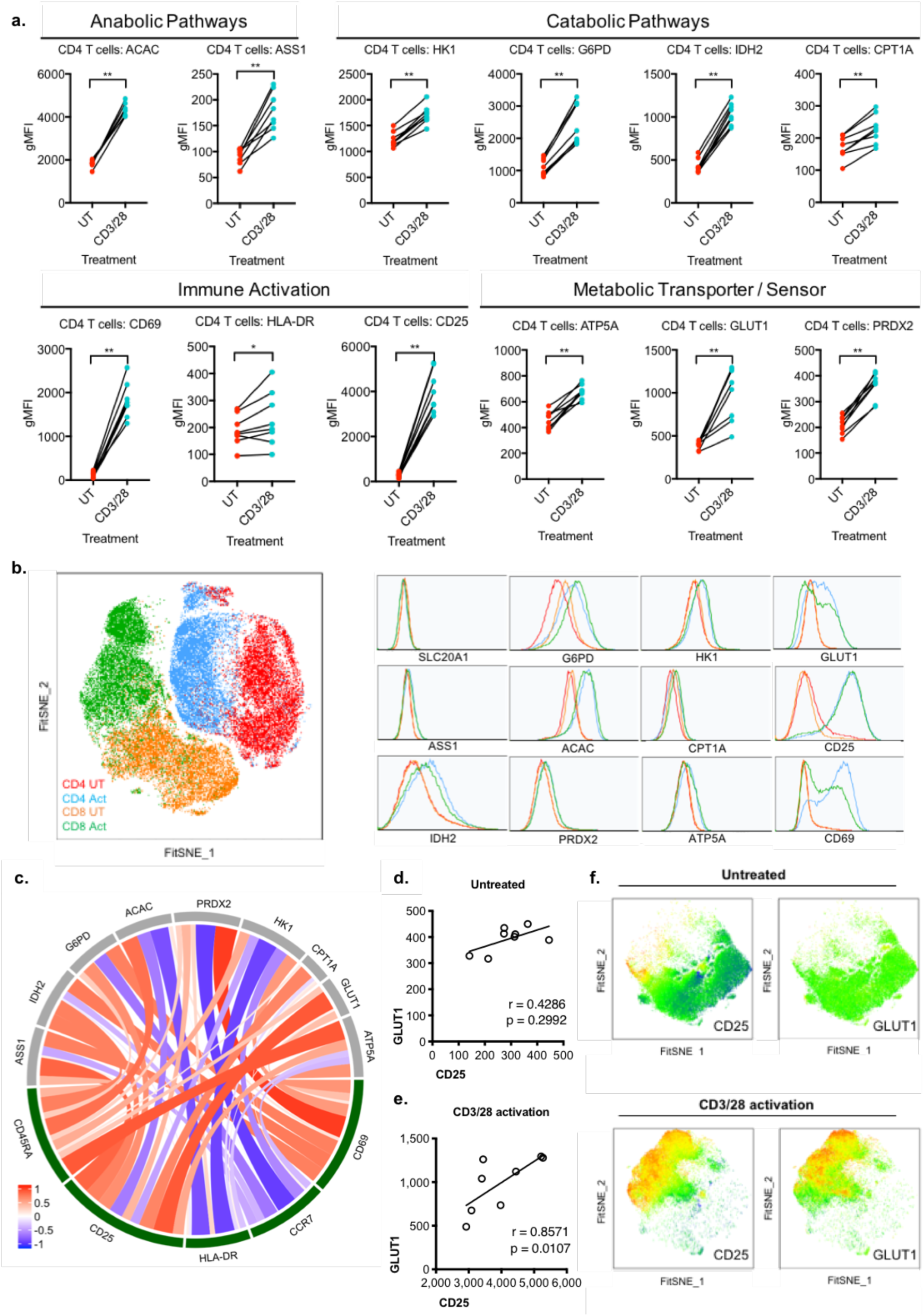
Activation induces extensive metabolic reprogramming in T cells. Purified T cells were untreated (UT) or activated with anti-CD3/CD28 beads (CD3/28). (a) Geometric mean fluorescence intensity (gMFI) was measured for activation and metabolic proteins in CD4^+^ T cells. Each dot represents one donor, data representative of n=8 donors, from 3 independent experiments. (b) FitSNE projection and corresponding expression of metabolic protein and activation markers in T cells, data acquired from n=5 samples, with 10,000 cells per donor. (c) Chord visualization of correlation between immune and metabolic proteins in activated CD4^+^ T cells, representative of n=8 donors. (d) Spearman correlation of GLUT1 and CD25 expression in untreated and (e) activated CD4^+^ T cells. (f) Heatmap of FitSNE projection of GLUT1 and CD25 expression in untreated and activated CD4^+^ T cells.

Congruent with the findings in purified T cells, the mixed PBMC studies showed similar increases in the capacity for flux through the glycolytic, PPP, OXPHOS and fatty acid metabolism pathways (Supplementary Fig. 3c). Whilst the purified T cells demonstrated a significant increase of ASS1 and PRDX2 protein level with activation, this observation was less pronounced in total PBMCs (Supplementary Fig. 3c). In contrast to the decreased expression of the phosphate transporter SLC20A1 with activation in PBMCs, we noted the loss of SLC20A1 once T cells were purified (Supplemental Fig. 3d). This was independent of any stimulation and suggests that purification methods alter the expression of some metabolic proteins.

We next investigated the relationship between metabolic state and T cell activation (Fig. 2c). The activation markers CD25 and CD69 showed positive correlations with multiple metabolic proteins. Conversely, a negative correlation between ACAC and HK1 with HLA-DR was demonstrated, indicating a difference in metabolic requirements of fatty acid synthesis and glycolysis for early and late activation. Specifically, the strongest correlation was seen between GLUT1 and CD25 (r=0.8571), indicating a positive relationship between the capacity for glucose uptake and CD25 expression with activation (Fig. 2d,e), and demonstrated by the overlap of high expressing CD25 and GLUT1 cells in the FitSNE projection of activated CD4^+^ T cells (Fig. 2f). This finding directly correlates the sensitivity of the T cell to the IL-2 growth factor CD25, to the capacity for glucose uptake by GLUT1, and is supported by the increase in glucose uptake by 2-NBDG (Supplementary Fig. 3e). In comparison to CD8^+^ T cells, we further demonstrate differential metabolic upregulation in CD4^+^ T cells with activation. Though at resting state, the CD4^+^ and CD8^+^ subsets show similar metabolic profiles, CD4^+^ T cells upregulate oxidative metabolism with higher expression of IDH2 and ATP5A, as well as GLUT1. In contrast, CD8^+^ T cells augment their capacity for flux through the PPP with higher G6PD expression (Fig. 2b, Supplementary Fig. 3e). This confirmed that T cell activation requires remodeling of the metabolic state that is specific to functional T cell subsets^29^.

Together, Met-Flow confirms previously described metabolic inductions of glycolysis, OXPHOS and fatty acid synthesis in activated T cells ^1,27–29^. This technique enabled the association of glycolysis and immune activation on a single cell level, by elucidating the positive correlation between GLUT1 and CD25 expression. The data further demonstrated reprogramming of other pathways, including key enzymes in mitochondrial respiration, the PPP and fatty acid oxidation, that contribute to the metabolic state of activated T cells.

### Glycolytic inhibition alters the metabolic and activation status of T cells

Previous analysis of global metabolic reprograming showed an increased capacity for glucose uptake and glycolysis, associated with T cell activation. Therefore, we investigated the dependence on glycolytic metabolism for the immuno-metabolic state. The glucose analog 2-Fluoro-2-deoxyglucose (FDG) was added in the presence or absence of anti-CD3/28 stimulation in purified T cells. 2-FDG is a closer analog to glucose than 2-DG, is less toxic, and more specific, as it does not interfere with mannose metabolism by incorporating into N-linked glycosylation^30–33^. We determined that 24 hours of 2-FDG alone did not cause a significant decrease in any metabolic pathway components (Fig. 3a-d). Stimulation of anti-CD3/28 resulted in an increase in surface expression of CD25 (Fig. 2a), whilst the addition of 2-FDG prevented this increase (Fig. 3c) ^34^. The dependence of glycolysis for CD25 expression was not shared across all surface activation molecules, since CD69 and HLA-DR were unchanged or increased respectively (Fig. 3c, Supplementary Fig. 4a). Inhibition of glycolysis with 2-FDG did not affect the protein level of GLUT1, indicating a feedback loop and the requirement for GLUT1 to maintain high levels of intracellular glucose (Fig. 3d, Supplementary Fig. 4a). As shown previously, stimulation with CD3/28 upregulated the expression of all other metabolic proteins analyzed, whilst 2-FDG combined with CD3/28 treatment reduced their expression with differential sensitivity, indicating partial dependence on glycolysis (Fig. 3a, b, d, Supplementary Fig. 4a). Taken together, these results indicate a heavy reliance on glucose for metabolic function during T cell activation.

**Figure 3.**
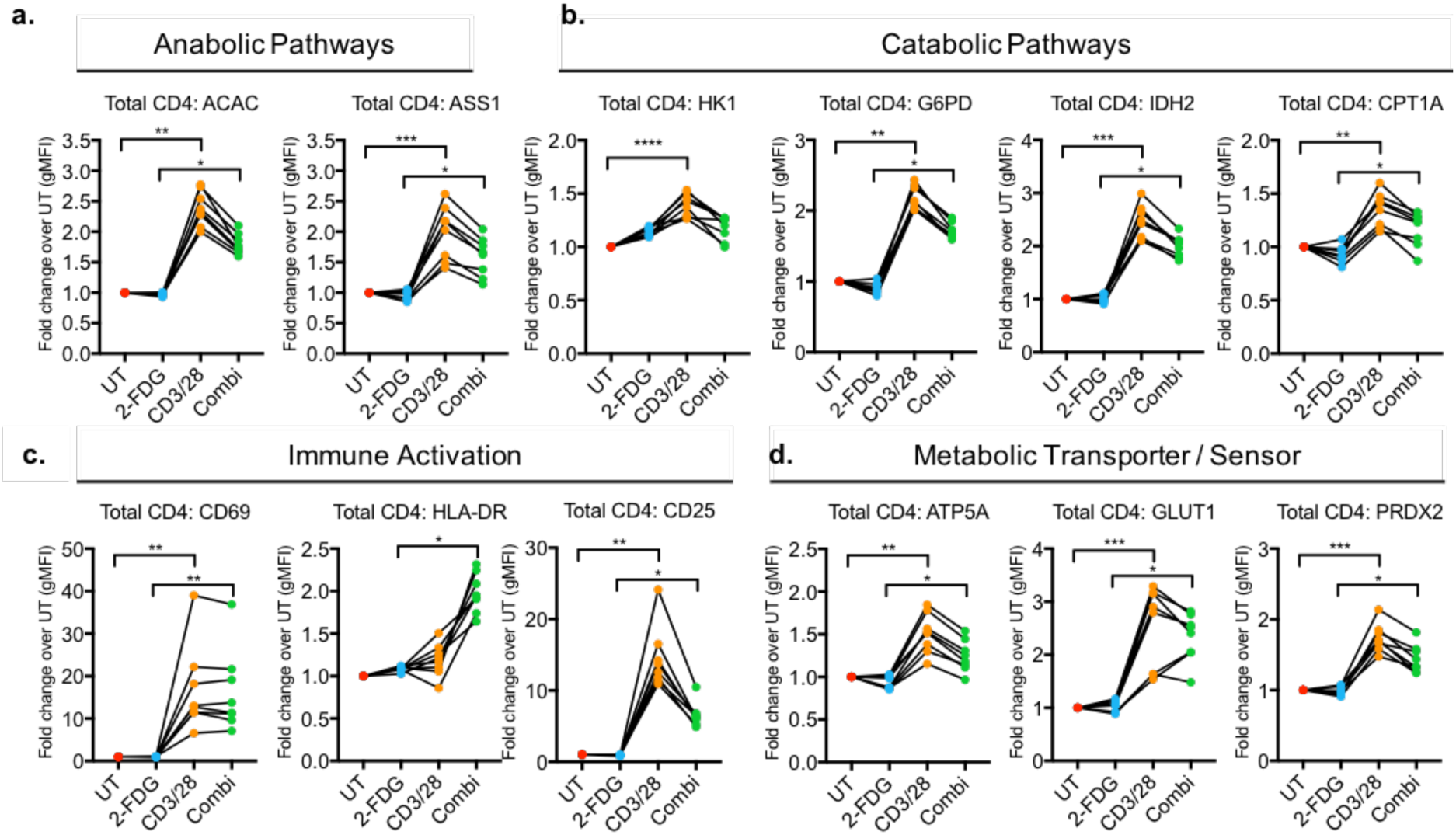
The activation and metabolic states of CD4+ T cells are altered by glycolytic inhibition. Fold change of metabolic protein and activation markers (gMFI) was measured in CD4^+^ T cells with no treatment (UT), 2-FDG, CD3/28, and combination of 2-FDG with CD3/28 (Combi). Metabolic proteins are grouped by (a) anabolic pathways, including fatty acid synthesis and arginine metabolism, and (b) catabolic pathways, including glycolysis, oxidative PPP, TCA cycle and fatty acid oxidation. (c) Activation markers and (d) the ATP synthase protein critical for OXPHOS, glucose transporter and the antioxidant protein were measured. Each dot represents one donor sample, total n=8 donors from 3 independent experiments.

To correlate the changes in maturation and metabolism of these T cells with cellular function, we measured the cytokine and chemokine release in their supernatants. This analysis showed a significant increase in pro-inflammatory CCL3, IL-13, IL-6, sCD40L, IL-17A, TNF-⍺, IFN-γ and CXCL10 following CD3/28 stimulation, as expected (Supplementary Fig. 4b). Glycolytic inhibition with 2-FDG selectively reduced the production of IL-13, IL-6, sCD40L, IL-17A. In contrast, IL-8 and GM-CSF increased following stimulation in the presence of 2-FDG, suggesting a regulatory role of glycolysis for these molecules (Supplementary Fig. 4b).

With the differential effects of glycolytic inhibition on activation markers and metabolic protein levels, our data demonstrated the dependence on glycolysis in regulating multiple metabolic pathways that alters T cell cytokine release. We showed glycolytic requirement for the upregulation of specific activation molecules and cytokines, including CD25, IL-13, IFN-γ and IL-17A. Moreover, all metabolic proteins were expressed at a lower level following glycolytic inhibition, with the exception of GLUT1, indicating maintenance of metabolic feedback. Collectively, Met-Flow is effective at elucidating differential responses of metabolic pathways in immunological processes.

### T cell memory subsets show differential metabolic phenotypes

In the studies described above, we showed the use of Met-Flow in assessing dynamic metabolic remodeling in T cell subsets following activation. Past studies have shown that T cell subsets utilize distinct energy sources under differential nutrient availability ^29,35–38^. Leveraging the capability of Met-Flow to measure metabolism of cellular subsets, we investigated the different metabolic states during memory differentiation and the effect of glycolytic inhibition.

Using FitSNE projection, the 10 metabolic proteins are differentially expression across memory subsets (Fig. 4a-b). We showed distinct sub-clusters of CM and EM populations based on their immuno-metabolic profiles, whereas the naïve and TEMRA subsets showed some overlap (Fig. 4a). The CM and EM populations both expressed higher levels of ACAC, PRDX2, and CPT1A, in contrast to the naïve and TEMRA subsets (Fig. 4b). Previous work has shown that EM cells have higher oxygen consumption rates and spare respiratory capacity in comparison to naïve CD4^+^ T cells ^2,35^. We corroborated this finding by showing increased IDH2 expression in the EM population (Fig. 4b, Supplementary Fig. 5a). Moreover, there is a concomitant high expression of PRDX2 in the EM cells, which may be a result of high oxidative stress produced by OXPHOS. These findings illustrate the ability to capture the differential metabolic states across resting T cell memory subsets using Met-Flow.

**Figure 4.**
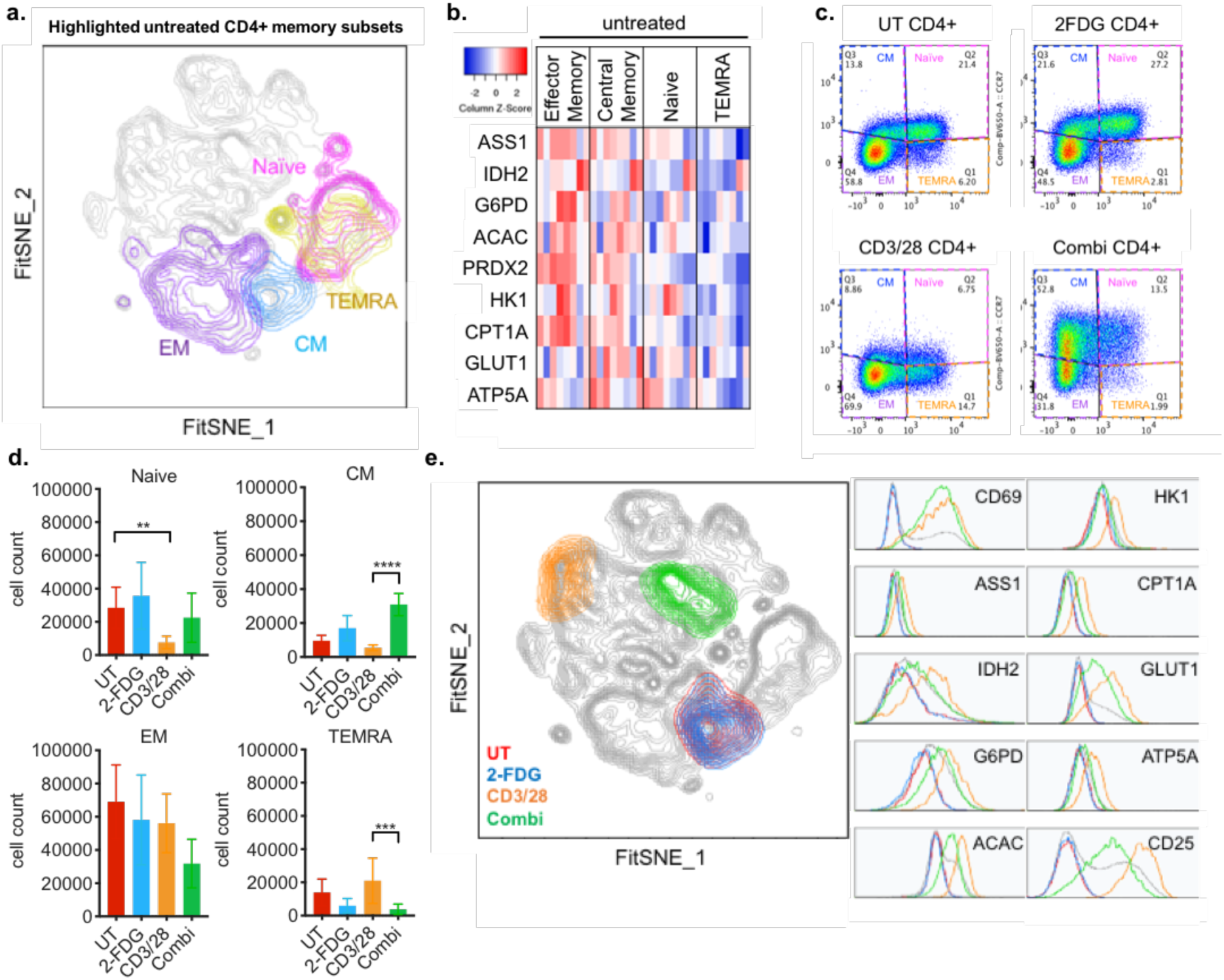
T cell memory subsets differentially respond to glycolytic inhibition. (a) FitSNE projection of resting state CD4^+^ memory populations, data represents n=5 donor samples. (b) Metabolic protein expression of resting state CD4^+^ memory subsets by gMFI, data represents n=8 donor samples. (c) Gating strategy of CD4^+^ memory subsets by CCR7 and CD45RA. (d) Cell count of CD4^+^ T memory populations across treatments. (e) FitSNE of CD4^+^ CM populations across treatments, data represents 5 donor samples from 2 independent experiments, with 20,000 cells per samples.

To measure the effect of glycolytic inhibition on the metabolic state across subsets, the cells were stimulated with CD3/28 and 2-FDG. This resulted in differential effects in each memory population, measured by cell frequency, metabolic protein level and activation status. Stimulation with CD3/28 caused a decrease in frequency of naïve CD4^+^ T cells compared to the untreated control (Fig. 4c-d). Addition of 2-FDG during activation (Combi) resulted in the increased frequency of CM cells and reduction of both TEMRA and EM populations (Fig. 4d). To further explore this expanded CM subset, we focused on the immuno-metabolic differences within the CM populations across treatment. The results demonstrated that glycolytic inhibition attenuated the activation induced expression of HK1, GLUT1, CPT1A, IDH2, G6PD, ACAC, ATP5A, PRDX2, ASS1 compared to activated CM cells (Fig. 4e, Supplementary Fig. 5b). This coincided with the decrease in CD25, but not in CD69 or HLA-DR, highlighting the difference in glycolytic dependence in early and late activation (Fig. 4e, Supplementary Fig. 5b). Lastly, compared to all other memory subsets and treatments, the FitSNE projection demonstrated a well-defined cluster based on the immuno-metabolic state of this perturbed CM subset (Fig. 4e, Supplementary Fig. 5c), indicating a specific metabolic state of this memory population.

Taken together, we show that Met-Flow is able to dissect metabolic profiles within T cell memory subsets. We identified the selective expansion of CM cells, that are independent of glycolysis, compared to other memory T subsets. Met-Flow captures divergent immuno-metabolic states in cellular subpopulations that arise during different cellular and tissue environments.

### Increased respiration and downstream pathway signaling in activated T cells

To confirm the metabolic reprogramming shown by flow cytometry, we assessed real-time respiration in bulk T cells using extracellular flux analysis, which analyses glycolytic function and mitochondrial respiration. We further identified a CM subset with high S6 phosphorylation, that expanded with 2-FDG addition. T cells were activated with CD3/28, and the corresponding metabolic modulators were added sequentially as described previously. As expected, the addition of CD3/28 induced a significant increase in overall glycolytic function, shown by elevated basal glycolysis, glycolytic capacity and reserve, compared to untreated cells (Fig. 5a,b). Mitochondrial respiration was also significantly impacted, revealing enhanced basal and maximal respiration, as well as spare mitochondrial capacity (Fig. 5c,d). These metabolic shifts in glycolysis and OXPHOS detected by extracellular flux analysis confirmed our metabolic protein flow cytometry results (Fig. 2b). Moreover, these changes in real-time respiration are supported by earlier work showing remodeling of glycolysis, the TCA cycle and OXPHOS following T cell activation ^29,39,40^.

**Figure 5.**
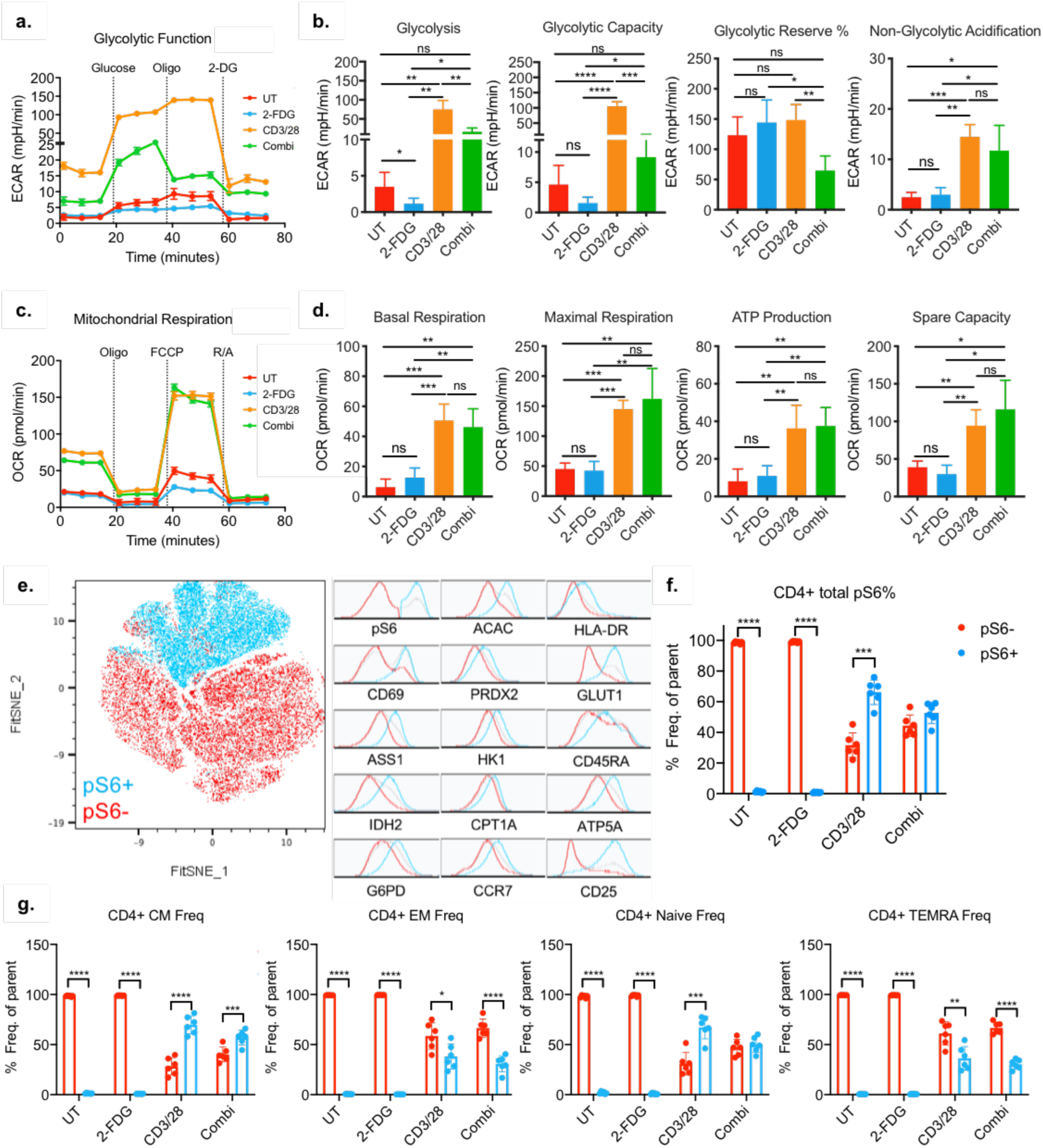
Respiration and downstream AKT signaling increase with T cell activation. (a) Glycolytic function across untreated (UT), 2-FDG treated, CD3/28 activated and combination treated (2-FDG+CD3/28) donor samples. Graph depicts one representative sample from a single donor. (b) Glycolytic parameters measured by extracellular acidification rate (ECAR) across treatments. (c) Mitochondrial respiration measured by oxygen consumption rate (OCR) in purified T cells across treatments and its associated (d) mitochondrial parameters. Data represents n=6 donor samples from 2 independent experiments. Statistical analysis was performed using one-way ANOVA with Tukey’s multiple comparisons test. (e) CD4+ T cells phosphorylation status of phospho-S6 (pS6) and respective levels of metabolic and activation markers. Data shown represents n=6 by FitSNE analysis. (f) Phosphorylation status across different treatments. (g) Phosphorylation status across memory subsets with treatment. Statistical analysis was performed using multiple t-test and Wilcoxon Signed Rank test.

We next evaluated the dependence of energetic metabolism on glucose using 2-FDG in real-time respiration. The activation-induced increases in ECAR and the associated glycolytic parameters were reduced in the presence of 2-FDG, confirming our earlier Met-Flow results (5a,b). Overall mitochondrial respiration did not significantly decrease with 2-FDG addition (Fig. 5c,d, Supplementary Fig. 6), indicating that at the bulk level their OXPHOS is not dependent on glucose.

Bulk analysis did not show a concurrent decrease in mitochondrial respiration with glycolytic inhibition, indicating cellular dependence on other carbon sources. We therefore aimed to investigate whether this dependence on alternative carbon sources was true for the entire population or specific for a subset of cells within bulk analysis. To evaluate the dynamics of metabolic protein level changes, we incorporated the phosphorylation state of ribosomal protein S6 (pS6) into the Met-Flow panel. The S6 protein is downstream of the mTORC1 signaling pathway and is phosphorylated upon TCR engagement, driving the translation of glycolytic proteins in T cells ^2,41^. Met-Flow analysis showed increased levels of CD69, CD25 and GLUT1 in the pS6 positive cells compared to pS6 negative cells, whereas the other metabolic proteins showed heterogeneous expression (Fig. 5e). This phosphorylation was specifically induced by CD3/28 stimulation, as untreated or 2-FDG treated T cells are pS6 negative. However, 2-FDG dampened this activation induced increase in the bulk population (Fig. 5f). To distinguish subset-specific metabolic preferences, we gated the T cell memory subsets using the expression of CCR7 and CD45RA to identify naïve, central memory (CM), effector memory (EM) and terminally differentiated effector memory T cells (TEMRA). Stimulation caused increased S6 phosphorylation across all memory subsets, compared to the unstimulated conditions. In the naïve and CM subsets, there was a mean of 67% and 69% pS6 positive cells respectively, whilst EM and TEMRA were 38% and 36%. The addition of 2-FDG to the stimulation condition caused the majority of cells to become pS6 negative in all subsets, apart from CD4 CM T-cells, in which the majority of cells maintained their pS6 positivity, thus, indicating their dependence their dependence on carbon sources other than glucose. These findings demonstrate the ability of Met-Flow to identify cellular populations with alternative metabolic reliance, which would not be achievable using other methodologies.

In sum, the bulk real-time respiration analysis confirms our previously described differential effect of activation and glycolytic inhibition on the metabolic state of T cells. Overall, increased downstream mTOR signaling corresponded with T cell activation in all memory subsets. We additionally identified a CM subset that was highly phosphorylated and glycolytically independent. Unlike bulk analysis, using Met-Flow identifies specific metabolic reprogramming corresponding to particular T cell subsets. These studies corroborate the changes in metabolic protein levels demonstrated by Met-Flow and further emphasize the unique advantages of single-cell metabolic flow cytometry over bulk analysis.

### Metabolic remodeling drives GM-CSF production of central memory T cells

We previously demonstrated that IL-8 and GM-CSF increased with glycolytic inhibition in bulk T cells, unlike other effector cytokines. To determine whether the metabolically distinct CM population was responsible for this inflammatory cytokine production, we measured the production of GM-CSF by incorporating a capture antibody into the Met-Flow capability. The production of GM-CSF by activated T cells stimulates myeloid cells to promote tissue inflammation^42,43^. Our data confirmed an increased production of GM-CSF with CD3/28 stimulation. This was linked to a higher metabolic state (Fig. 6a), whereas unstimulated or only 2-FDG treated T cells produce low GM-CSF and showed lower levels of metabolic protein expression (Fig. 6a, Supplementary Fig. 7a). We next investigated whether GM-CSF production was different across T cell memory subsets. With CD3/28 treatment, the EM subset was the largest GM-CSF producing population (Fig. 6b). The addition of 2-FDG showed a selective reduction of GM-CSF production in the EM and TEMRA memory populations. In contrast, the CM subset increased with 2-FDG addition to CD3/28, and the naïve population showed a similar trend (Fig. 6b-c). This increase in the GM-CSF producing CM cells was similar to the expanded pS6 high CM population (Fig. 4g), demonstrating glycolytic independence specific for this memory subset.

**Figure 6.**
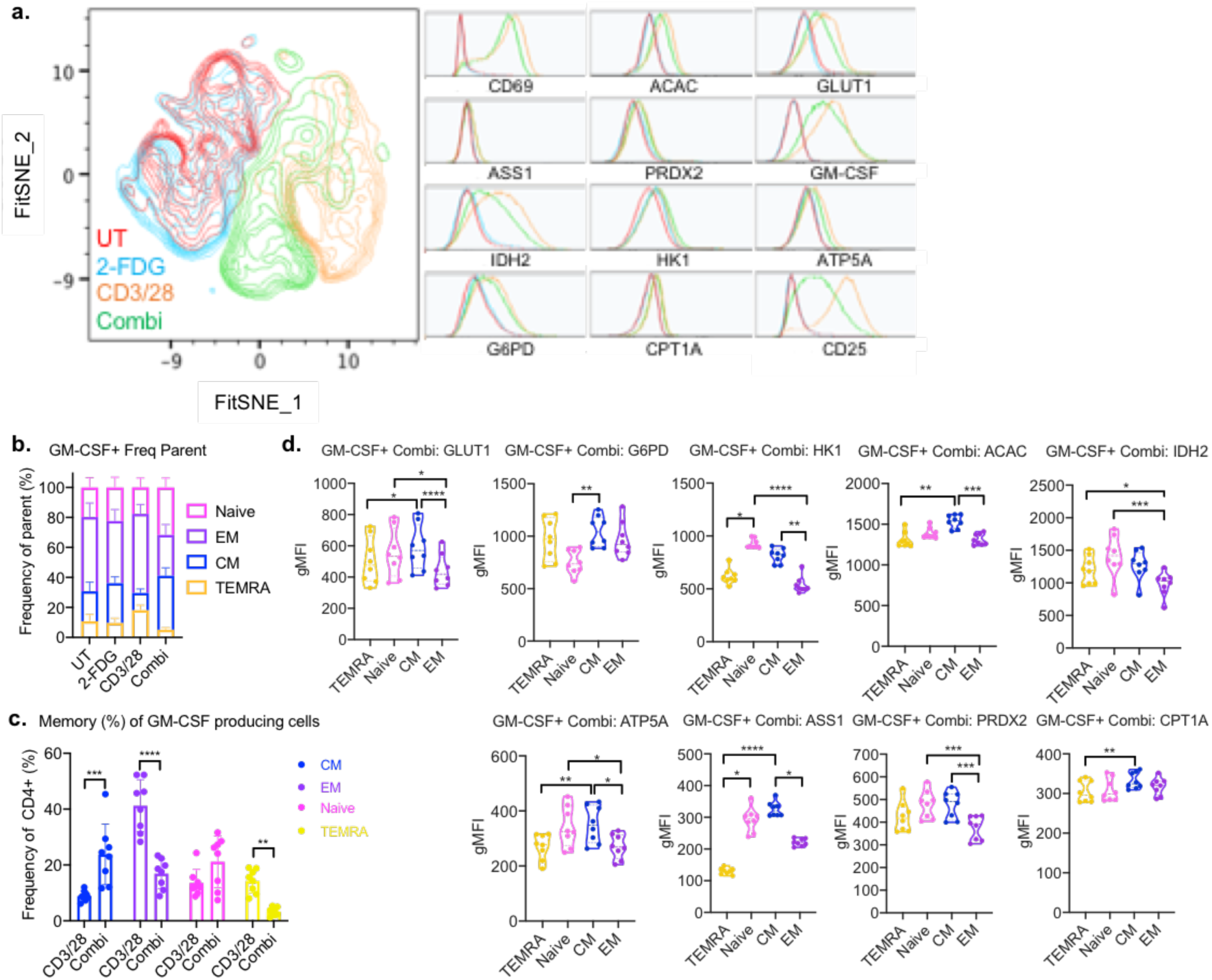
Glucose restriction and metabolic remodeling drive the expansion of inflammatory memory T subpopulation. (a) FitSNE projection of GM-CSF producing total CD4+ T cells. (b) GM-CSF producing frequency (%) of CD4+ T cells across all treatments and memory subsets. (c) Comparison of activation and combi (CD3/28+2-FDG) treated memory subset frequency. (d) Differential expression of metabolic proteins across T cell memory subsets with glycolytic inhibition during CD3/28 activation (combi). All data represents n=8 donors, and statistical analysis was performed using T test or Friedman’s test with Dunn’s multiple comparisons.

To link the differential GM-CSF producing subsets to their underlying metabolic state, we measured their metabolic protein expression. The decrease in GM-CSF production with 2-FDG addition in EM was associated with lower metabolic protein expression compared to CD3/28 treatment alone (Supplementary Fig. 7b). Comparing across subsets with combination activation and glycolytic inhibition, the CM subsets shows an overall trend of high metabolic protein expression (Fig. 6d). Specifically, in comparison to the glycolytically dependent EM subset, the CM population was characterized by higher expression of glycolytic proteins, GLUT1 and HK1, as well as increased fatty acid synthesis enzyme ACAC, OXPHOS protein ATP5A, arginine synthesis by ASS1 and the antioxidant enzyme PRDX2. Unlike other memory subsets, this increased frequency of GM-CSF producing CM population has a specific metabolic state, that is differentially impacted by glycolytic inhibition.

In conclusion, the expansion of Met-Flow with cytokine analysis demonstrated the ability to attribute differential effector function to divergent metabolic states of specific immune subsets. Using this capability, we identified a novel metabolic profile of pro-inflammatory CM T cells, which produce high GM-CSF independently of glucose metabolism.

## Discussion

The ability to measure the metabolic state of specific immune cells is essential for a fundamental understanding of cellular function, particularly in the context of disease. Here, we present Met-Flow, a novel capability to simultaneously measure multiple metabolic pathways across diverse immune subsets on a single-cell, protein level using a combination of intracellular staining and flow cytometry. The application of this technology on human PBMCs revealed cell type specific differences in core metabolic pathways. Furthermore, we demonstrate that the surface expression of specific activation molecules, cytokines, and chemokines were dependent on their underlying metabolic state in a cell type specific manner. Together, this novel technique demonstrates that immune cell subsets have unique metabolic protein signatures relating directly to their activation and maturation states.

Bulk cellular analysis has demonstrated that leukocytes possess an array of metabolic states leading to different functional capacity and disease outcome^35,44^. Moreover, the metabolic microenvironment and tissue localization influence immune cell function ^45–47^. We began by analyzing the total peripheral blood mononuclear population using Met-Flow, enabling a global view of immune cell metabolism. Analysis of monocytes, which play a key role in innate immune function, revealed a higher expression of all metabolic proteins relative to other cell types. This suggests that they exist in a metabolically poised state with implications for inflammatory responses and plasticity^48,49^. This elevated expression was not due to assay intrinsic factors such as cellular size, as the mDC and monocyte populations share a similar forward scatter profile, yet their metabolic protein expression is vastly different (data not shown). Deeper subset analysis demonstrated that the inflammatory CD86^+^CD16^+^ monocytes expressed higher HK1, suggesting a greater glycolytic capacity. This supports and clarifies earlier work showing that the activation induced upregulation in the global monocyte population is dependent on glycolysis ^50^. In addition our findings show divergent metabolic requirements in the DC subpopulations following activation ^51^. We observed different metabolic profiles, with a higher capacity for flux through arginine metabolism in mDCs, compared to higher capacity for OXPHOS and glucose uptake in pDCs. The characterization of NK subsets using Met-Flow revealed that CD56 bright cells express significantly higher HK1, confirming their increased glycolytic activity in comparison to the CD56 dim subset^52^. Moreover, we highlight an opposing requirement for fatty acids, with CD56 bright cells expressing higher fatty acid synthesis enzyme ACAC, compared to the increased capacity for flux through fatty acid oxidation by CPT1A in the CD56 dim cells. NKT cells had higher levels of IDH2 in comparison to CD4^+^ T cells, which verifies studies that show increased levels of OXPHOS in NKT cells, important for their function ^53^. Taken together, this demonstrates the ability of Met-Flow to simultaneously analyze diverse metabolic states on differential immune subpopulations.

The association of immune and metabolic states has been extensively studied in T cell biology. Increased utilization of glycolysis, OXPHOS, and fatty acid synthesis following T cell receptor stimulation ^1,34,39,54–56^. In this study, we confirmed these findings and further demonstrated the involvement of the PPP, fatty acid oxidation, antioxidant level, and arginine synthesis pathways post activation. Using our method, we confirmed the highly oxidative phenotype of CD4^+^ in comparison to the CD8^+^ subset^29^. Though expression of the glycolytic enzyme HK1 was similar between both subsets, the PPP was significantly induced in CD8^+^ T cells, indicating a differential metabolic program that utilizes glucose breakdown. We also confirmed that the activation induced expression of the high affinity IL-2 receptor, CD25, is dependent on glycolysis. Importantly, CD25 expression positively correlated with GLUT1 protein levels, confirming the association with activation induced glucose uptake. Similar associations between GLUT1 and CD25 expression were found in activated CD8+ T cells from chronic lymphocytic leukemia (CLL) patients ^57^. These CLL patient-derived T cells showed lower GLUT1 intracellular reserves upon stimulation, and impaired mitochondrial fitness compared to activated T cells from healthy donors. This highlights the potential of Met-Flow to measure reprogramming of immuno-metabolic states in the diseased context. Studies have shown that CD8^+^ memory cells have a higher mitochondrial capacity and favor fatty acid oxidation compared to the naïve counterparts ^58,59^. Moreover, the inhibition of the glycolytic pathway enhances the formation of CD8^+^ memory cells ^60,61^. By leveraging the single-cell nature of the technology, we were able to dissect T cell memory populations into EM and CM based on the combination of surface markers and intracellular metabolic profiles and found differential regulation by glycolysis. At resting state, our data confirms higher metabolic activity in CM and EM subsets compared to naïve T cells ^2^. With the inhibition of glycolysis, we observed an expansion of the CM population, whereas the frequency of the EM population decreased. A lower reliance on fatty acid synthesis was previously shown in CD4^+^ EM cells in low glucose conditions, whereas CM and naïve populations can increase their uptake of fatty acids for survival ^38^. Our data similarly demonstrates that the metabolic state of CM cells was distinct from other memory populations both with and without glycolytic inhibition. This illustrates the ability to capture differential responses of cellular subpopulations by revealing diverse immuno-metabolic states, reflecting divergent metabolic dependence and function.

Consistent with previous studies, the release of cytokines post activation was largely dependent on glycolysis^1^. We showed that glycolytic inhibition decreased IL-13, IL-6, sCD40L and IL-17A production from isolated T cells. Interestingly, GM-CSF and IL-8 were not dependent on glycolysis suggesting differential control and redundancy in the metabolic regulation of cytokine production. A rapid immune response, including the production of cytokines, is regulated by multiple post-transcriptional mechanisms, including non-coding RNAs, micro-RNA, RNA-binding proteins and translational control by mTOR signaling ^13,62–64^. Specifically, GM-SCF mRNA stability is controlled by protein binding to AU-rich elements in 3’-untranslated regions, which direct mRNA degradation and control mRNA half-life^65,66^. The intersection of cytokine biology and metabolism is often regulated at the post-transcriptional level. Similarly to GM-CSF, the production of IFNγ and TNFα are controlled by the repression of mRNA binding of lactate dehydrogenase ^37,67^. Like many genes involved in dynamic processes, post-transcriptional regulation of cytokine production can make accurate measurement of gene expression using mRNA abundance difficult^64^.

GM-CSF production by T cells activates myeloid cells for inflammatory cytokine production, phagocytosis and pathogen killing^68–70^. Pro-inflammatory T cells are known to be associated with negative disease outcome, as GM-CSF production can drive disease progression in autoimmune disorders^71^, neurological disease^72^ and skin hyperinflammation^73^. In graft-versus-host disease, high GM-CSF produced by allogeneic T cells induces donor-derived myeloid cells to produce inflammatory cytokines, which drives pathology^74^. In hepatocellular carcinoma, tumor cells produce high amounts of GM-CSF that recruit myeloid derived suppressor cells to induce immune tolerance and increase PD-L1 expression^75^. Using Met-Flow analysis, we identified the selective expansion a CM subset with a unique metabolic state, that produces high amounts of GM-CSF in the absence of glucose. This population expressed high GLUT1, ACAC, PRDX2, ATP5A, ASS1 and HK1, indicating a metabolic state independent of glycolysis. This profile was specific to the CM subset, as the activated EM cells reduced total metabolic activity and frequency of GM-CSF producing cells with glycolytic inhibition. Using Met-Flow, we have discovered a novel metabolic phenotype of a clinically important T cell subset. This suggests an axis of pro-inflammatory T cell differentiation relevant in multiple inflammatory pathologies. Inhibiting GM-CSF production by targeted restriction of the metabolic pathways identified using Met-Flow could give rise to novel therapeutic targets for combination with tumor immunotherapy.

In summary, the studies presented here have described a novel high dimensional flow cytometry technique, which facilitates the analysis of key metabolic proteins, cellular lineage and activation molecules simultaneously. Traditional methods assess metabolism in bulk populations, which do not have the ability to identify differences in the metabolic profile in cellular subsets on a single cell, protein level. These methods can mask important attributes specific to infrequent populations and may not account for heterogeneity in cellular subsets. Using our method, we were able to simultaneously capture dynamic metabolic states across multiple immune populations. Met-Flow can be combined with methods of post-translational modification such as histone acetylation, phosphorylation status and intracellular cytokine production, enabling comprehensive single-cell immuno-metabolic analysis at the protein level. The expansion of this technique with the inclusion of additional biosynthetic pathways, will be greatly assisted with improvements in other high-dimensional flow based methods, such as Abseq^76^ and Cytof^77^. Met-Flow can be applied to the investigation of metabolic remodeling in any cell type and disease context and has the potential to uncover unique metabolic targets for therapeutic intervention.

## Acknowledgments

We sincerely thank Sriram Narayanan and Marius Jones for their assistance with reading the manuscript and for their helpful advice. We would also like to thank Robert Balderas, Keefe Chee, William Wong, and John Wotherspoon from Becton Dickinson and Company for their contribution in custom antibody conjugation and flow cytometry panel design. This study was supported by the Agency for Science, Technology and Research Singapore and the National Medical Research Council Singapore (NMRC/OFLCG/003/2018).

## Author contributions

P.J.A. and J.E.C. designed and directed the study. P.J.A., N.K., R.H. and B.A. setup and performed experiments. W.X. performed computational analysis. P.J.A., B.A., W.X., R.H., N.K. and J.E.C. analyzed and interpreted the data. P.J.A., R.H., A.M.F, and J.E.C. wrote the manuscript.

