## Supplementary Figures for "A novel strategy for single-cell metabolic analysis highlights dynamic changes in immune subpopulations"

| Gene Name | Gene description | Metabolic pathway | Function (HPA, RefSeq) | Localization (HPA) |
|---|---|---|---|---|
| *SLC20A1/ PiT1* | Solute carrier family 20 member 1 | Phosphate transporter | Sodium-dependent phosphate import, promotes apoptosis and regulates Akt-1 [51] | Vesicles |
| *ASS1* | Argininosuccinate Synthase 1 | Arginine biosynthesis | Rate-limiting step of urea cycle, de novo arginine synthesis, alters p53-AKT signalling [52] | Cytosol, Nucleoplasm |
| *SLC2A1/ GLUT1* | Solute carrier family 2 member 1/ facilitated glucose transporter member 1 | Glucose uptake | Glucose import, response to hypoxia and glucose starvation | Plasma Membrane |
| *IDH2* | Isocitrate dehydrogenase (NADP(+)) 2 | TCA cycle | NADP+ dependent, NAPDH producing, role in energy production [53] | Mitochondria |
| *G6PD* | Glucose-6-phosphate dehydrogenase | Oxidative PPP | Rate limiting step of oxidative PPP, provides NADPH and pentose phosphates for fatty acid and nucleic acid synthesis[54] | Cytosol, MTOC, vesicles |
| *ACAC/ ACC1* | Acetyl-CoA Carboxylase Alpha | Fatty acid synthesis | Rate limiting step in de novo long chain fatty acid synthesis | Cytosol, nucleoli fibrillar centre |
| *PRDX2* | Peroxiredoxin 2 | Antioxidant | Peroxiredoxin family of antioxidant enzymes, prevents oxidative stress | Cytosol |
| *HK1* | Hexokinase 1 | Glycolysis | Rate-limiting enzyme, catalyses first step of glucose metabolism, couples glycolysis to intramitochondrial OXPHOS | Mitochondria |
| *CPT1A* | Carnitine palmitoyl-transferase 1A | Fatty Acid Oxidation | Key enzyme in carnitine-dependent transport across the mitochondrial inner membrane | Outer Mitochondrial Membrane |
| *ATP5A/ ATP5A1/ ATP5F1A* | ATP synthase F1 subunit alpha | ATP biosynthesis | Catalyses ATP synthesis, ATP-binding soluble catalytic core, H+ transport | Mitochondria |

**Supplementary Table 1. Metabolic proteins representing critical components and rate-limiting enzymes of metabolic pathways.** Metabolic proteins and their function are based on transcriptomic and proteomic analysis reported in the Human Protein Atlas (HPA) [15,16] and by Reference sequence database at NCBI (RefSeq) [17].

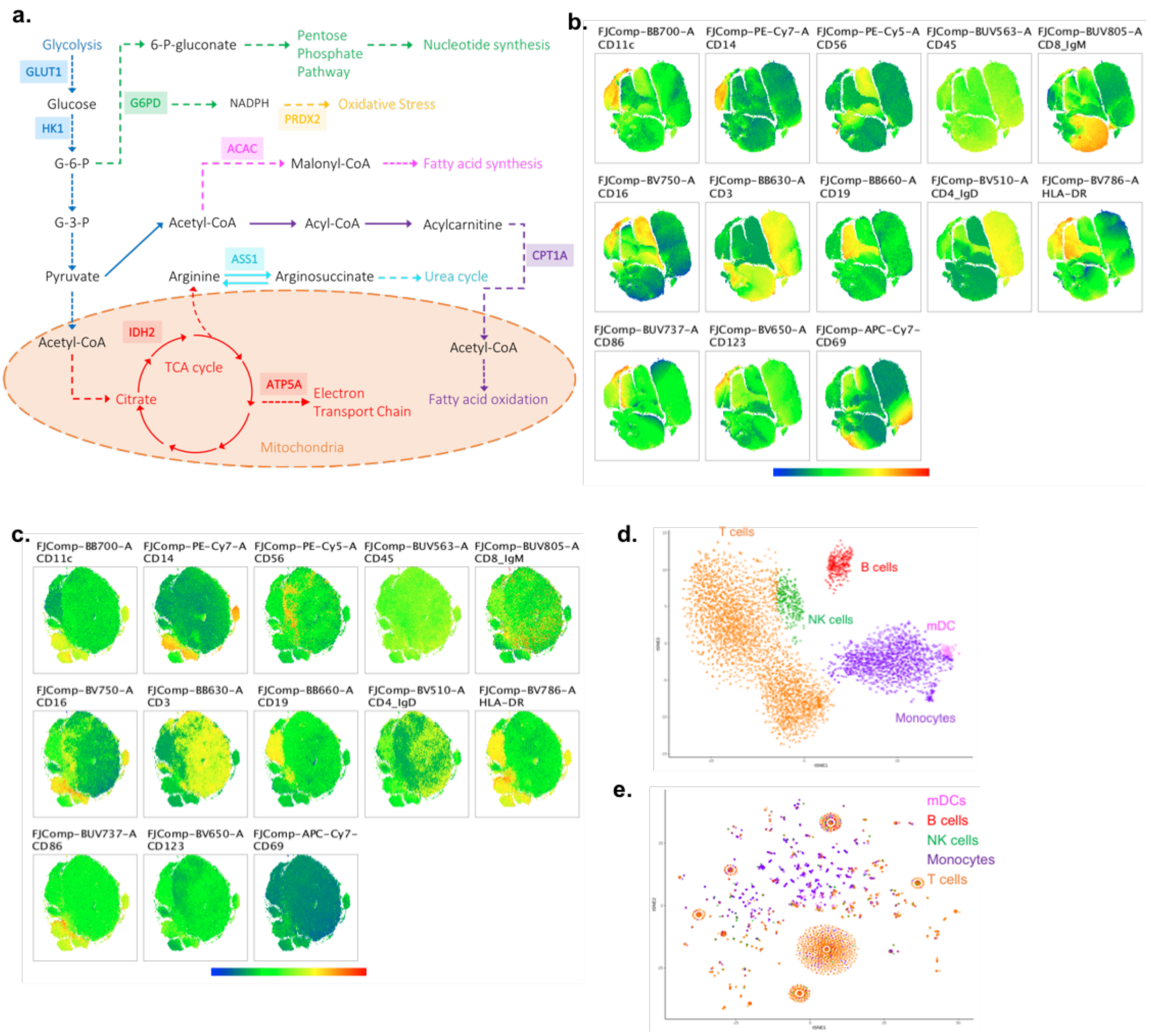


**Supplemental Figure 1. Metabolic pathways and proteins analyzed using Flow Cytometry.** (a) Ten specific metabolic enzymes were selected, which are critical and rate-limiting in each distinct pathway. These represent glucose metabolism by the glucose transporter (GLUT1), Hexokinase 1 (HK1), the pentose phosphate pathway by Glucose-6-Phosphate Dehydrogenase (G6PD), oxidative stress regulation by Peroxiredoxin 2 (PRDX2), phosphate import by Solute carrier family 20 member 1 (SLC20A1), fatty acid metabolism by Acetyl-Co-A Carboxylase Alpha (ACAC) and Carnitine-Palmitoyl Transferase 1A (CPT1A), urea cycle by Arginosuccinate-Synthetase 1 (ASS1), and oxidative phosphorylation by mitochondrial isocitrate dehydrogenase (IDH2) and ATP synthase F1 subunit alpha (ATP5A). (b) Heatmap of phenotypic markers to identify immune populations on the FitSNE projection of immune markers with metabolic protein level, (c) and on the projection of metabolic proteins only. (d) tSNE projection of scRNAseq in purified PBMC using >500 metabolic genes, and (e) the same scRNAseq dataset using the corresponding RNA levels of the 10 Met-Flow metabolic proteins.

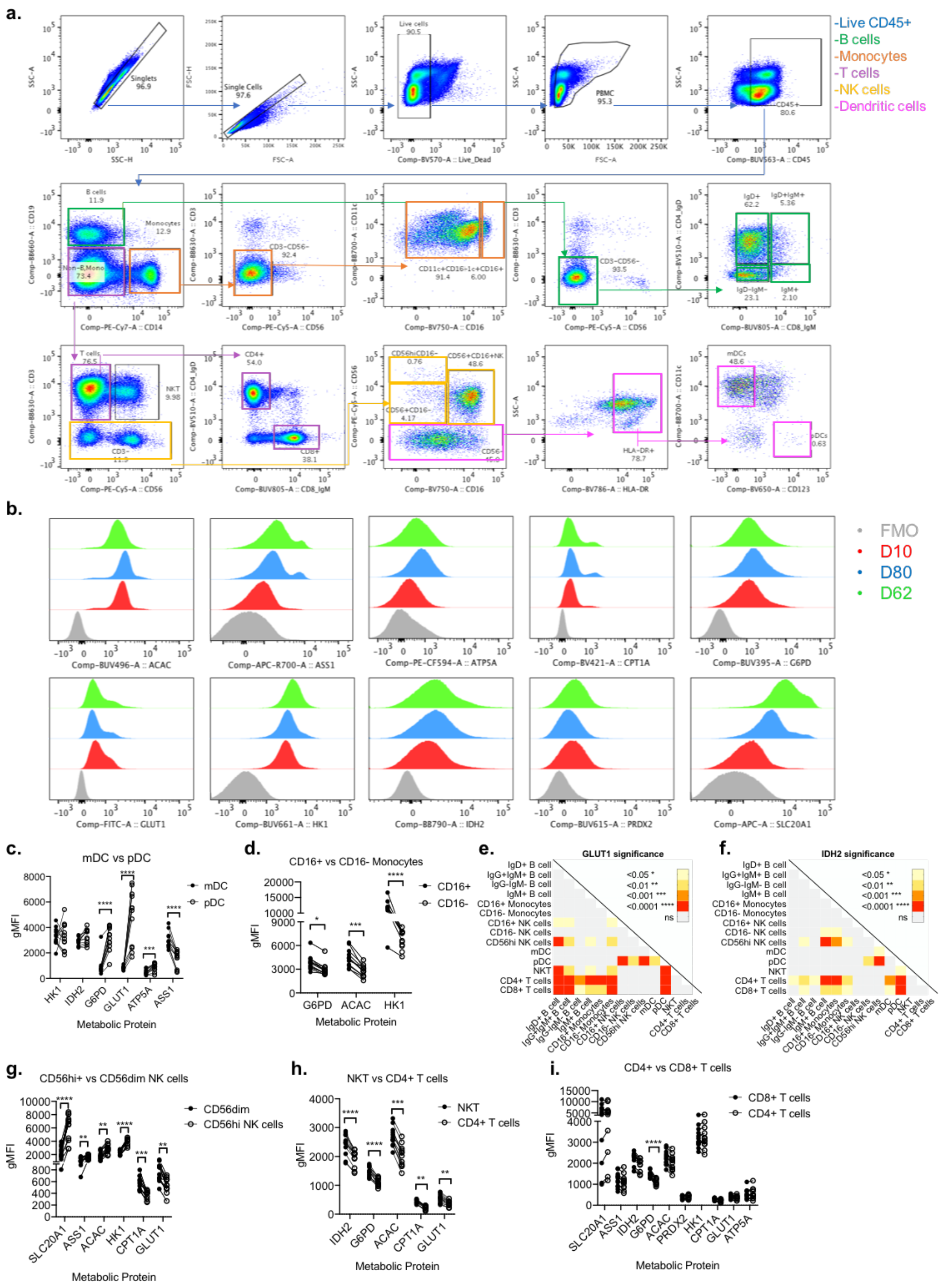


**Supplementary Figure 2. Divergent metabolic protein levels across immune subsets.** (a) Gating strategy for each immune population. (b) Expression of each metabolic protein in 3 healthy donors stained with all antibodies, compared to the fluorescence-minus-one (FMO) control, stained with all except the corresponding antibody. (c) Selected metabolic target expression between mDC and pDCs, (d) and CD16$^+$ and CD16$^-$ Monocytes. (e) Expression of GLUT1 (f) and IDH2. Colors indicate significant p values measured by One-way ANOVA with Friedman multiple comparisons test. (f) Selected metabolic target expression between CD56$^{hi}$ and CD56$^{dim}$ (CD56$^+$CD16$^-$), (h) NKT and CD4$^+$ T cells, and (i) between CD8$^+$ and CD4$^+$ T cells, statistical significance measured by multiple T-test with Holm-Sidak multiple comparisons.

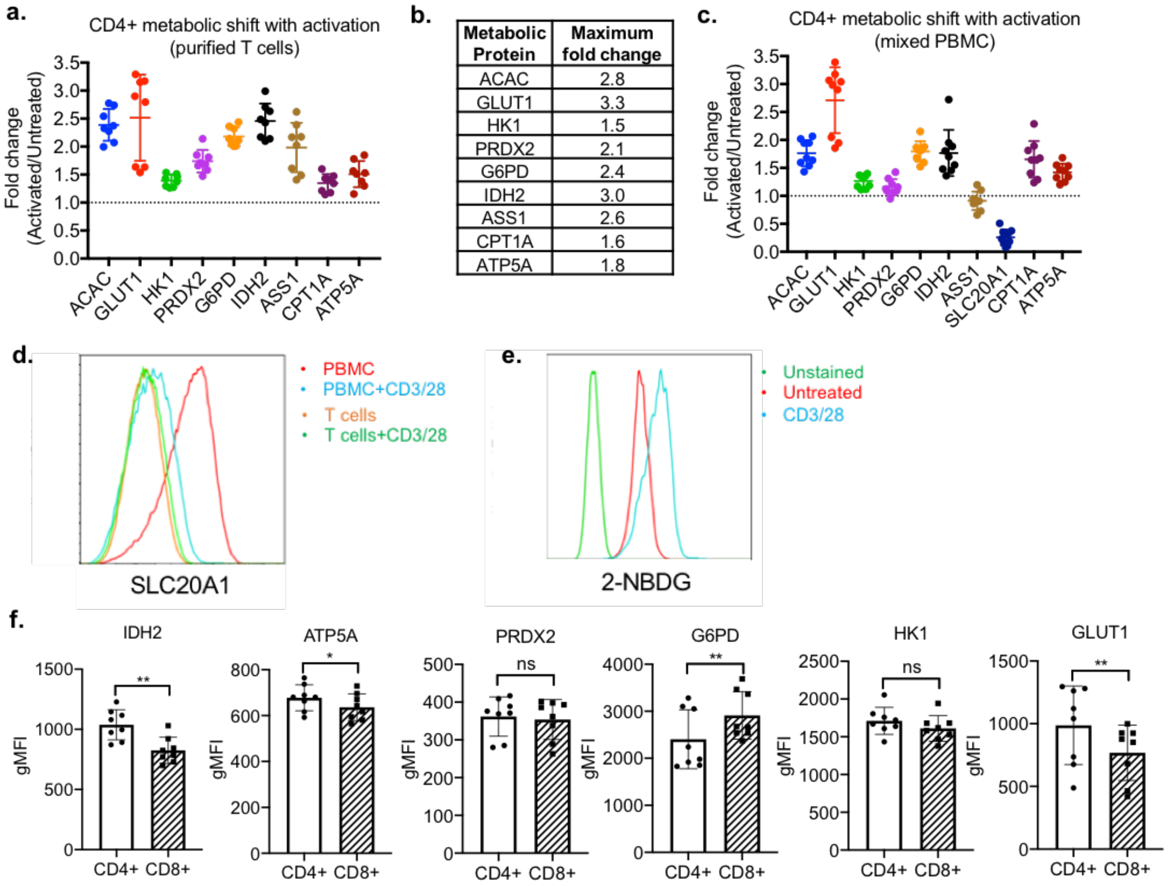


| Metabolic Protein | Maximum fold change |
|---|---|
| ACAC | 2.8 |
| GLUT1 | 3.3 |
| HK1 | 1.5 |
| PRDX2 | 2.1 |
| G6PD | 2.4 |
| IDH2 | 3.0 |
| ASS1 | 2.6 |
| CPT1A | 1.6 |
| ATP5A | 1.8 |

**Supplementary Figure 3. Comparison of individual metabolic proteins in CD4+ T cells**
(a) Metabolic shift with activation, calculated by fold change of expression by activated over untreated gMFI in CD4+ gated in purified T cells and (b) with maximum fold change values. (c) Expression of metabolic targets in CD4+ T cells from a mixed PBMC population. (d) Expression of SLC20A1 in CD4+ T cells in a mixed sample of PBMC or purified T cells using the same flow cytometry panel. (e) Glucose uptake measurement by 2-NBDG (200uM) in purified T cells by flow cytometry. (f) Comparison of oxidative and glycolysis associated proteins between CD4+ and CD8+ isolated T cells. Data represents n=8 donors, using paired T-test, Wilcoxon matched signed ranked test.

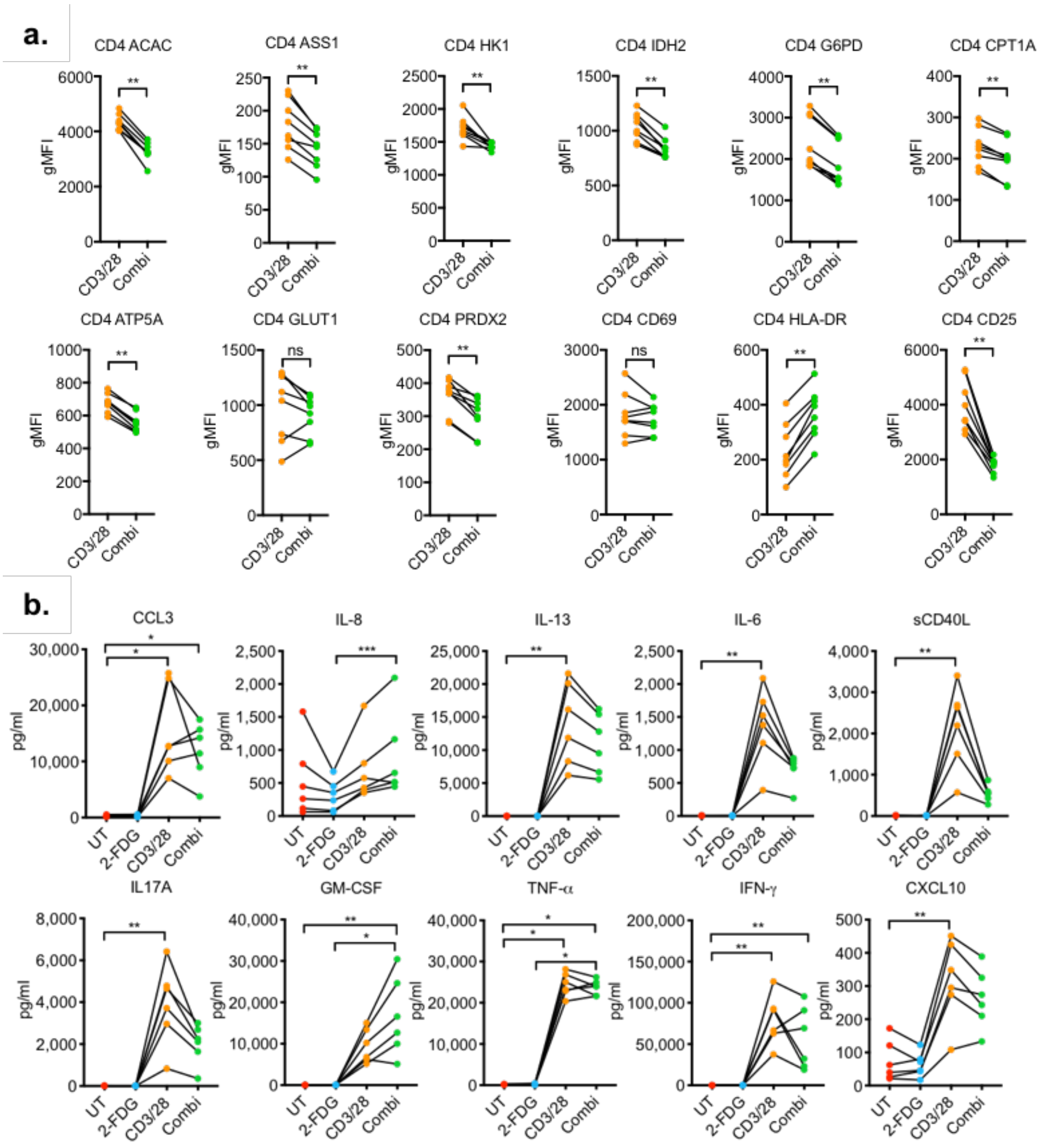


**Supplementary Figure 4. Effect of combination (2-FDG+CD3/28) treatment on T cells** (a) Expression of metabolic targets between activation (CD3/28) vs. combination (2-FDG+CD3/28) treatment, using paired T-test, Wilcoxon matched signed ranked for analysis. (b) Cytokine and chemokine production in untreated T cells, 2-FDG alone, CD3/28 alone, Combi (2-FDG+CD3/28). Experiments represent n=6 donors in 2 independent experiments.

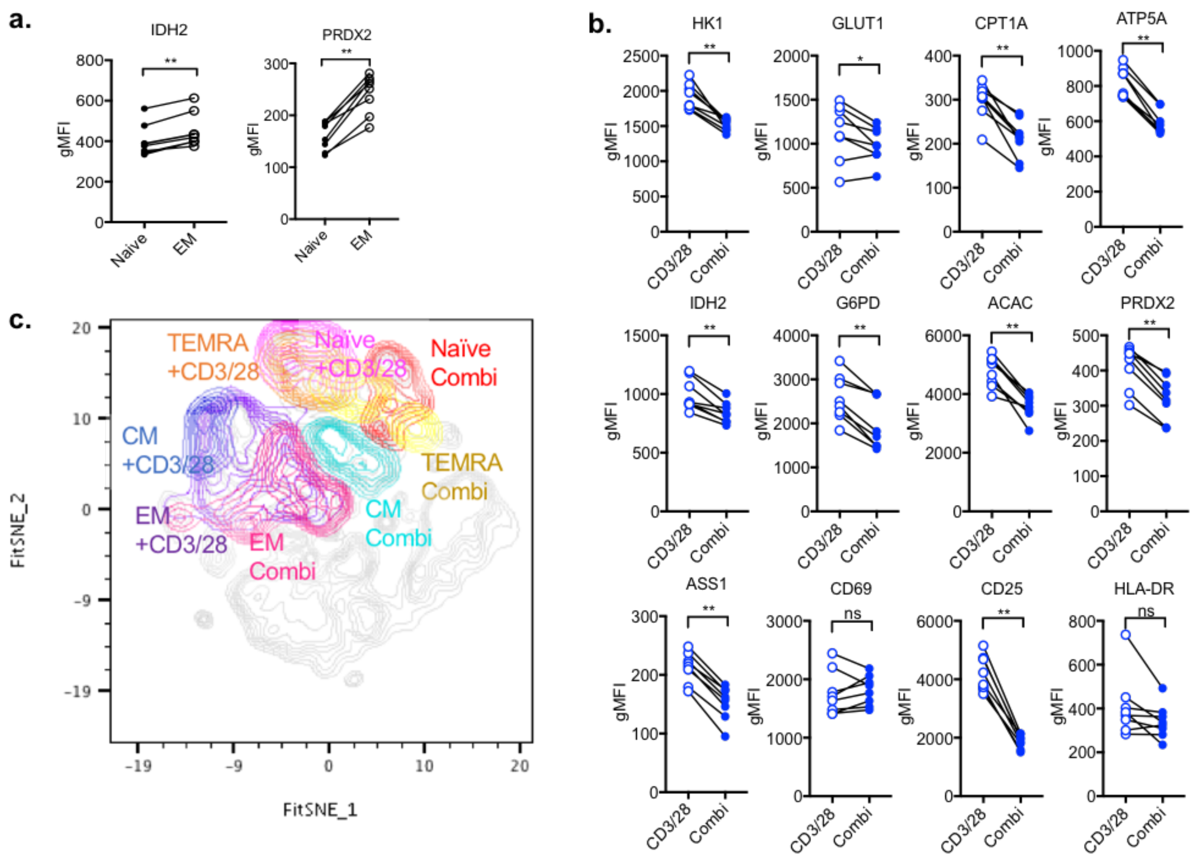


**Supplementary Figure 5. Differential expression of metabolic targets across memory populations** (a) Comparison of Naïve vs EM at steady state. (b) Expression level of memory subsets with combi (CD3/28 + 2-FDG) treatment. (c) FitSNE projection of CD3/28 activated and combi treated memory subsets. Data represents n=5 samples.

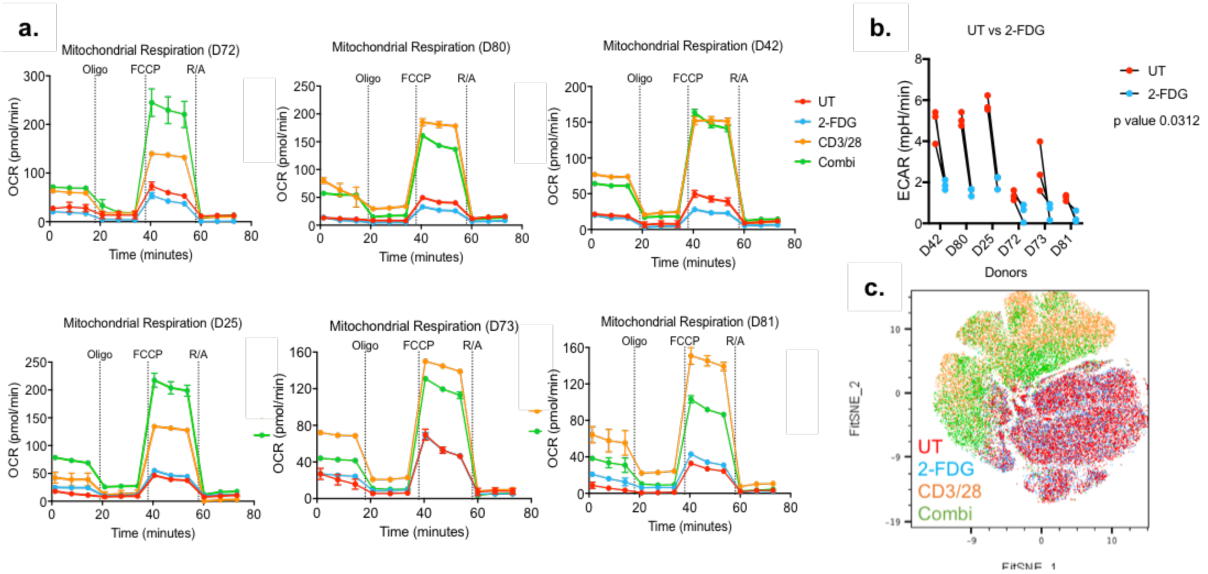


**Supplementary Figure 6. Extracellular flux analysis of combination (2-FDG+CD3/28) treatment on T cells** (a) Differential mitochondrial respiration rates across treatments in 6 donors. (b) Extracellular acidification rate (ECAR) between untreated (UT) and 2-FDG. (c) CD4+ T cells clusters based on treatment. Data shown represents n=6 by FitSNE analysis.

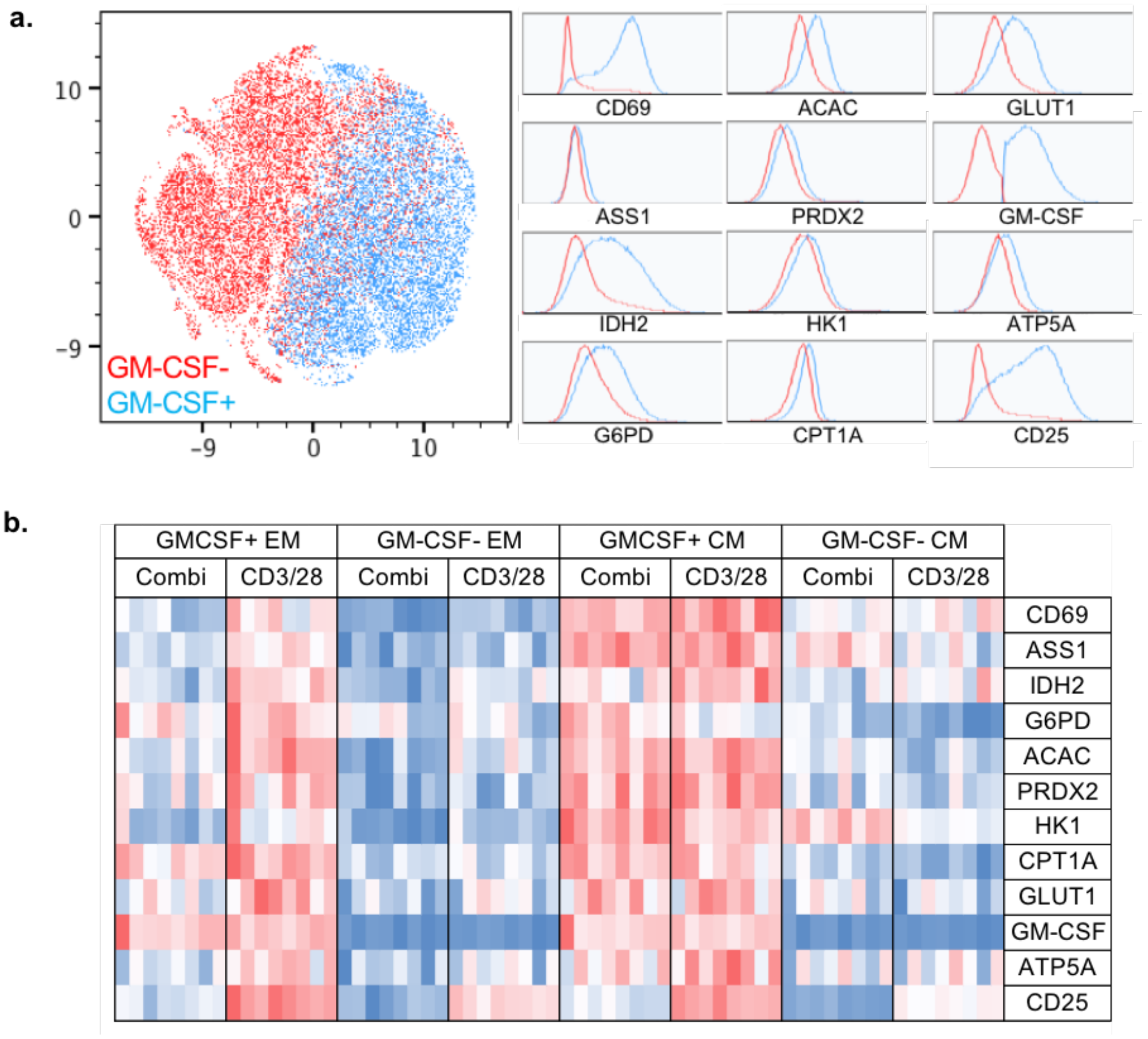


**Supplementary Figure 7. Differential metabolic profiles of GM-CSF producing memory T cells** (a) Metabolic state of GM-CSF+ vs GM-CSF- cells. (b) Expression of metabolic targets within the CM population, between activated (CD3/28) and combination (CD3/28+2-FDG) treatment, calculated by non-parametric paired T-test, Wilcoxon matched pairs signed-rank test.
